# *Lmx1b* influences correct post-mitotic coding of mesodiencephalic dopaminergic neurons

**DOI:** 10.1101/441915

**Authors:** Iris Wever, Pablo Largo Barrientos, Elisa J. Hoekstra, Marten P. Smidt

## Abstract

The Lim Homeobox transcription factor 1 beta (LMX1b) has been identified as one of the transcription factors important for the development of mesodiencephalic dopaminergic (mdDA) neurons. During early development, *Lmx1b* is essential for induction and maintenance of the Isthmic Organizer (IsO), and genetic ablation results in the disruption of inductive activity from the IsO and loss of properly differentiated mdDA neurons.

To study the downstream targets of Lmx1b without affecting the IsO, we generated a conditional model in which *Lmx1b* was selectively deleted in *Pitx3* expressing cells from embryonic day (E)13 onward. Supporting previous data, no significant changes could be observed in general dopamine (DA) marks, like *Th, Pitx3 and Vmat2* at E14.5. However, in depth analysis by means of RNA-sequencing revealed that *Lmx1b* is important for the expression level of survival factors *En1* and *En2* and for the repression of mdDA subset mark *Ahd2* during (late) development. Interestingly, the regulation of *Ahd2* by *Lmx1b* was found to be *Pitx3* independent, since *Pitx3* levels were not altered in *Lmx1b* conditional knock-outs (cKO) and *Ahd2* expression was also up-regulated in *Lmx1b/Pitx3* double mutants compared to *Pitx3* mutants. Further analysis of *Lmx1b* cKOs showed that post-mitotic deletion of *Lmx1b* additional leads to a loss of TH+ cells at 3 months age both in the VTA and SNc. Remarkably, different cell types were affected in the SNc and the VTA. While TH+AHD2+ cells were lost the SNc, TH+AHD2- neurons were affected in the VTA, reflected by a loss of *Cck* expression, indicating that *Lmx1b* is important for the survival of a sub-group of mdDA neurons.

## Introduction

Dopamine (DA) is one of the catecholaminergic neurotransmitters found in the central nervous system. Although cell bodies of DA neurons can be found in several positions within the mammalian brain, the largest population of DA neurons is located in the ventral mesencephalon (Björklund and Dunnett, 2007; Roeper, 2013; Smidt and Burbach, 2007). The mesodiencephalic dopaminergic (mdDA) neuronal population consists of specific subsets with distinct functions (Smits et al., 2006), including the Substantia Nigra pars compacta (SNc) and the Ventral tegmental area (VTA). The SNc innervates the dorsolateral striatum and caudate putamen forming the nigrostriatal pathway. The nigrostriatal pathway is integrated in a complex network that controls voluntary movement and body posture, and the degeneration of this pathway is the characteristic of Parkinson’s disease (Braak et al., 2003). The VTA has projections to the ventral striatum, the amygdala and the prefrontal cortex. These pathways are involved in regulating emotion-related behavior and are linked to addiction, depression and schizophrenia (Prakash and Wurst, 2006; Smidt and Burbach, 2007). The different subsets of mdDA neurons can already be distinguished during embryonic development based on their molecular profile and anatomical position (La Manno et al., 2016; Smits et al., 2013). It has been shown that the rostrolateral population will eventually form the largest part of the SNc, while the mediocaudal population is destined to become the VTA (Hökfelt et al., 1980; Veenvliet et al., 2013). Recent studies have revealed that each subset is dependent on a different transcriptional program for their development and survival (Jacobs et al., 2011; Kouwenhoven et al., 2017; Panman et al., 2014; Smits et al., 2006; Veenvliet and Smidt, 2014; Veenvliet et al., 2013). The rostrolateral fate is determined by a complex interplay between *Pitx3* and *En1. En1* induces the general DA phenotype and influences *Pitx3* expression, while *Pitx3* is required for the suppression of the caudomedial phenotype by down-regulating *En1* levels ((Veenvliet et al., 2013). After the formation of the rostrolateral population the remaining DA population will obtain a caudomedial fate under the control of *En1* (Bye et al., 2012; Veenvliet et al., 2013). Another factor that has been shown to be crucial for mdDA development and neuronal survival is LMX1B (Adams et al., 2000; Deng et al., 2011; Doucet-Beaupré et al., 2016; Laguna et al., 2015a; Smidt et al., 2000). Analysis of the *Lmx1b* complete knock out mouse revealed that LMX1B is an essential component of a positive feedback loop required to maintain genes associated with the formation and functioning of the Isthmic organizer (IsO), including *Wnt1*, *En1*, *En2*, *Pax2* and *Fgf8* (Adams et al., 2000; Guo et al., 2007). In the absence of *Lmx1b* the initiation, maintenance and inductive activity of the IsO were found to be severely impaired, affecting the development of the midbrain in general (Guo et al., 2007). Further analysis of *Lmx1b -/-* embryonic midbrains demonstrated a reduction in TH-expressing cells in the ventral mesencephalon (Deng et al., 2011; Smidt et al., 2000). Although the loss of *Lmx1b* initially seemed to primarily affect the lateral group of the DA progenitor domain (Deng et al., 2011), the medially located TH-expressing cells failed to induce *Pitx3* and were lost during further development (Smidt et al., 2000). In contrast to the *Lmx1b* null mutant, the conditional deletion of *Lmx1b* in mdDA progenitors, but not in the IsO, resulted in normal development of mdDA neurons (Yan et al., 2011). It was suggested that *Lmx1a* compensates for the loss of *Lmx1b*, since the deletion of both led to a severe reduction in both DA progenitors and mature mdDA neurons (Yan et al., 2011). Further studies into the functional redundancy between *Lmx1a* and *Lmx1b* suggest that the function of *Lmx1b* in mdDA development and postnatal neuronal survival can be mostly compensated by *Lmx1a* (Deng et al., 2011; Doucet-Beaupré et al., 2016; Nakatani et al., 2010; Yan et al., 2011). However, a recent study by Laguna *et al.* identified a critical role for *Lmx1b* in the functioning of mdDA neurons. They showed that *DatCre* driven deletion of *Lmx1b* reduces the levels of DAT and TH in the nerve terminals of mdDA neurons in the dorsal and ventral Striatum of 18 months old mutant mice (Laguna et al., 2015). In the present study we aimed to further elucidate the role of *Lmx1b* in postmitotic development and neuronal survival of mdDA neurons. We generated a mouse model that deletes *Lmx1b* in postmitotic mdDA neurons, by crossing a floxed *Lmx1b* mutant (Suleiman et al., 2007) with a *Pitx3* driven *iCre* (Smidt et al., 2012). The loss of *Lmx1b* did not affect the development of the general DA phenotype, however a loss of TH+ cells was observed in 3 month old animals, suggesting that *Lmx1b* plays a role in the survival and/or maintenance of mdDA neurons. During (late) development we found reduced levels of *En1* and *En2*, which relates to the function of *Lmx1b* during early development (Guo et al., 2007). In addition, the rostrolateral subset mark, *Ahd2*, was increased and ectopically expressed at E14.5, however, the caudomedial mark *Cck* was unaffected. To verify whether mdDA subsets were also affected during adult stages, the amount of TH+AHD2+ cells were analyzed in both the SNc and the VTA. Interestingly the amount of TH+AHD2+ cells was less in the SNc of the mutant compared to controls, but not in the VTA. In the VTA, TH+AHD2- cells were affected, which was reflected in lower levels of *Cck.* Taken together, our data shows that *Lmx1b* plays a role during the late development and survival of mdDA neurons. During development *Lmx1b* is important for maintaining proper *En1* and *En2* levels and the repression of the subset mark *Ahd2*, while during adult stages *Lmx1b* is important for the survival of a sub-group of mdDA neurons.

## Material and Methods

### Ethics Statement

All animal studies are performed in accordance with local animal welfare regulations, as this project has been approved by the animal experimental committee (Dier ethische commissie,Universiteit van Amsterdam; DEC-UvA), and international guidelines.

### Animals

All lines were maintained on a C57BL/6J background (Charles River). *Lmx1b*-floxed animals were generated by R.L. Johnson and obtained from R. Witzgall (University of Regensburg, Germany) and have been previously described (Suleiman et al., 2007). The *Pitx3Cre* has been previously generated in our lab (Smidt et al., 2012). *Pitx3CreCre; Lmx1b* L/+ animals were crossed with *Lmx1b* L/+ or *Pitx3CreCre; Lmx1b* L/+ animals to generate *Pitx3Cre/+; Lmx1b* +/+, *Pitx3Cre/+; Lmx1b* L/L and *Pitx3CreCre; Lmx1b* +/+, *Pitx3CreCre; Lmx1b* L/+ littermates. Embryos were isolated at embryonic day (E) 14.5, considering the morning of plug formation as E0.5. Pregnant and adult mice were euthanized by CO2 asphyxitian and embryos and brain were collected in 1xPBS and immediately frozen on dry-ice (fresh frozen) or fixed by immersion in 4% paraformaldehyde (PFA) for 4-8hrs at 4°C. After PFA incubation, samples were cryoprotected O/N in 30% sucrose at 4°C. Embryos and brains were frozen on dry-ice and stored at −80°C. Cryosections were slices at 16μm, mounted at Superfrost plus slides, air-dried and stored at −80°C until further use.

### Genotyping

Genotyping of the Lmx1b-Lox animals was done by PCR using primers: forward 5′- AGGCTCCATCCATTCTTCTC, and reverse: 5′-CCACAATAAGCAAGAGGCAC, resulting in a wild-type product of 243 bp, or a LoxP-inserted product of 277 bp. Pitx3-Cre genotyping was done by two different PCR’s. To discriminate between wild-type and Pitx3-Cre: forward 5′- GCATGATTTCAGGGATGGAC and reverse 5′-ATGCTCCTGTCTGTGTGCAG, resulting in a product of 750 bp for a mutant allele, and no product in wild-type animals. To additionally discriminate between heterozygous and homozygous Pitx3-Cre animals, primers were designed around Pitx3 exon 2 and exon 3: forward 5′-CAAGGGGCAGGAGCACA and reverse 5′-GTGAGGTTGGTCCACACCG, resulting in a product of 390 bp for the wildtype allele and no product for the mutant allele.

### In situ hybridization

In situ hybridization with digoxigenin (DIG)-labeled RNA probes was performed as described (Smidt et al., 2004; Smits et al., 2003). DIG-Labeled RNA probes for *Th, Vmat2, Dat, Aadc, Pitx3, En1, Ahd2* and *Cck* have been previously described (Grima et al., 1985; Hoekstra et al., 2013; Jacobs et al., 2007; Smidt et al., 2004; Smits et al., 2003). The used Lmx1b probe is a 310bp fragment containing exon 4-6 of the *Lmx1b* coding sequence.

### Fluorescence Immunohistochemistry

Cryosections were blocked with 4% HIFCS in 1x THZT [50mM Tris-HCL, pH 7.6; 0.5M NaCl; 0.5% Triton X-100] and incubated with a primary antibody [Rb-TH (Pelfreeze 1:1000), Sh-TH (Millipore AB1542, 1:750), Rb-AHD2 (Abcam AB24343, 1:500)] in 1xTHZT overnight at room temperature. The following day the slides were washed and incubated for 2 hrs at room temperature with secondary Alexafluor antibody (anti-rabbit, anti-sheep (In vitrogen, 1:1000) in 1xTBS. After immunostaining nuclei were staining with DAPI (1:3000) and embedded in Fluorsave (Calbiogen) and analyzed with the use of a fluorescent microscope (Leica). All washes were done in TBS and double stainings were performed sequential, with immunostaining for TH being done first, followed by the staining for AHD2. The antibody against AHD2 requires antigen retrieval, which was executed as follows; slides were incubates 10 minutes in PFA after which they were submersed in 0.1M citrate buffer pH 6.0 for 3 minutes at 1000Watts followed by 9 minutes at 200Watts. Slides were left to cool down, after which protocol was followed as described above.

### Quantitative analysis

Quantification of TH-expressing neurons, TH+AHD2+ cells and TH+AHD- cells in 3 month old and 12 month old midbrain was performed in ImageJ as follows. First TH-positive cells were counted independently of whether or not they expressed AHD2. Cells were counted in 10-12 matching coronal sections (3 months old, n=3 wildtype, n=4 conditional knock-out (cKO), 12 months old, n=4 wildtype, n=4 cKO). Cells were counted as TH+neurons when TH staining co-localized with nuclear DAPI staining. For the double stained sections a color overlay was made and the double stained cells were counted as TH+AHD2+ and the green cells were considered as TH+AHD2- cells. Similar to the single TH countings, cells were counted as neurons when the staining co-localized with nuclear DAPI staining. The separation of the SNc and VTA were made based on anatomical landmarks. Everything rostral of the supramammillary decussation was considered as SNc and distinction between the SNc and the VTA was made based on the tracts medial lemniscus, positioned between the SNc and VTA. Statistical analysis was performed via a student’s T-test.

### RNA-sequencing

RNA was isolated from dissected E14.5 midbrains of *Pitx3Cre/+; Lmx1b* +/+ and *Lmx1b* cKO embryos. RNA was isolated with Trizol (ThermoFisher) according to manufacturers protocol. After isolation RNA clean-up was performed with an RNA mini kit from Qiagen according to manufacturers protocol. RNA isolated of 2 wildtype or 2 mutant embryos was pooled per samples to eventually form an n=3. Pair-end RNA-sequencing (minimal 2* 10^7 reads per sample),mapping on the mouse genome and statistical analysis on read counts was performed by Service XS (Leiden, The Netherlands). The mapping was done with the mouse ensemble GRCm38.p4 data set.

### Quantitative PCR (qPCR)

RNA was isolated from dissected E14.5 midbrains of *Pitx3Cre/+; Lmx1b* +/+ and *Lmx1b* cKO embryos and of E14.5 *Pitx3CreCre; Lmx1b* +/+ and *Pitx3CreCre; Lmx1b* L/L. RNA was isolated with Trizol (ThermoFisher) according to manufacturers protocol. For the *Lmx1b* cKO 3 midbrains were pooled per samples (n=4 wildtype, n=4 cKO) and further purified on a column, according to the manufacturers protocol (Qiagen, RNeasy mini kit). For the *Pitx3/Lmx1b* mutants a single midbrain was used per sample (n=3 *Pitx3CreCre; Lmx1b* +/+, n=3 *Pitx3CreCre; Lmx1b* L/L) and no RNA clean-up was performed. Relative expression levels were determined by using the QuantiTect SYBR green PCR lightCycler kit (Qiagen) according to the manufacturers instructions. For each reaction 10ng (dissected midbrain) of total RNA was used as input. Primers used for *Th, Ahd2, En1* and *Cck* were previously published (Jacobs et al., 2011), primers for *Lmx1b* forward 5’-GAGCAAAGATGAAGAAGCTGGC and reverse 5’-CTCCATGCGGCTTGACAGAA (product size 98bp).

## Results

### Specific deletion of *Lmx1b* in developing *Pitx3* positive neurons

Studies on *Lmx1b* null mutants have shown that *Lmx1b* is essential for the development of mdDA neurons. In the absence of *Lmx1b* TH+ cells fail to induce *Pitx3* and are lost during development(Smidt et al., 2000). In addition to an effect on the mdDA neuron population, *Lmx1b* was also found to be essential for the expression of several genes associated with IsO functioning, including *Fgf8, Wnt1,En1, En2, Pax2* and *Gbx2*, and that the loss of *Lmx1b* severely affects the initiation, maintenance and the inductive activity of the IsO during midbrain development(Guo et al., 2007). To study the role of *Lmx1b* in mdDA development, without affecting the IsO, we generated a conditional *Lmx1b* knock-out. *Lmx1b*-floxed animals(Suleiman et al., 2007) were crossed with *Pitx3Cre* animals(Smidt et al., 2012), to generate mice that lack *Lmx1b* exclusively in *Pitx3* positive neurons. *Pitx3* expression is initiated around E11.5(Smidt et al., 1997) and it has been shown that the onset of CRE activity of *Pitx3*-Locus driven *Cre* expression is around E13.5(Smidt et al., 2012), so to guarantee a complete loss of *Lmx1b* transcript we decided to study E14.5 embryos. To verify that these animals lack *Lmx1b* only in the *Pitx3* positive domain, we assessed the expression of *Lmx1b* in *Pitx3Cre/+; Lmx1b L/L* compared to *Pitx3Cre/+;Lmx1b +/+* littermates by means of *in situ* hybridization (Figure 1A). A clear loss of *Lmx1b* expression could be observed in the mdDA area (arrowheads) in *Lmx1b L/L* embryos, while expression in other regions, like the diencephalon and the hindbrain, remains unaffected (arrows). To examine whether the loss of a *Pitx3* allele and the complete loss of *Lmx1b* influenced *Pitx3* expression we analyzed its expression in adjacent slides (Figure 1B). When comparing wild-type to mutant embryos no clear difference in *Pitx3* expression was observed. Taken together, our analysis shows that crossing *Lmx1b*-floxed mice with *Pitx3Cre* animals, generates a *Lmx1b* conditional knock-out, in which the deletion is confined to the mdDA neuronal field, leaving the rostral and caudal expression domains of *Lmx1b* intact.

**Figure 1:**
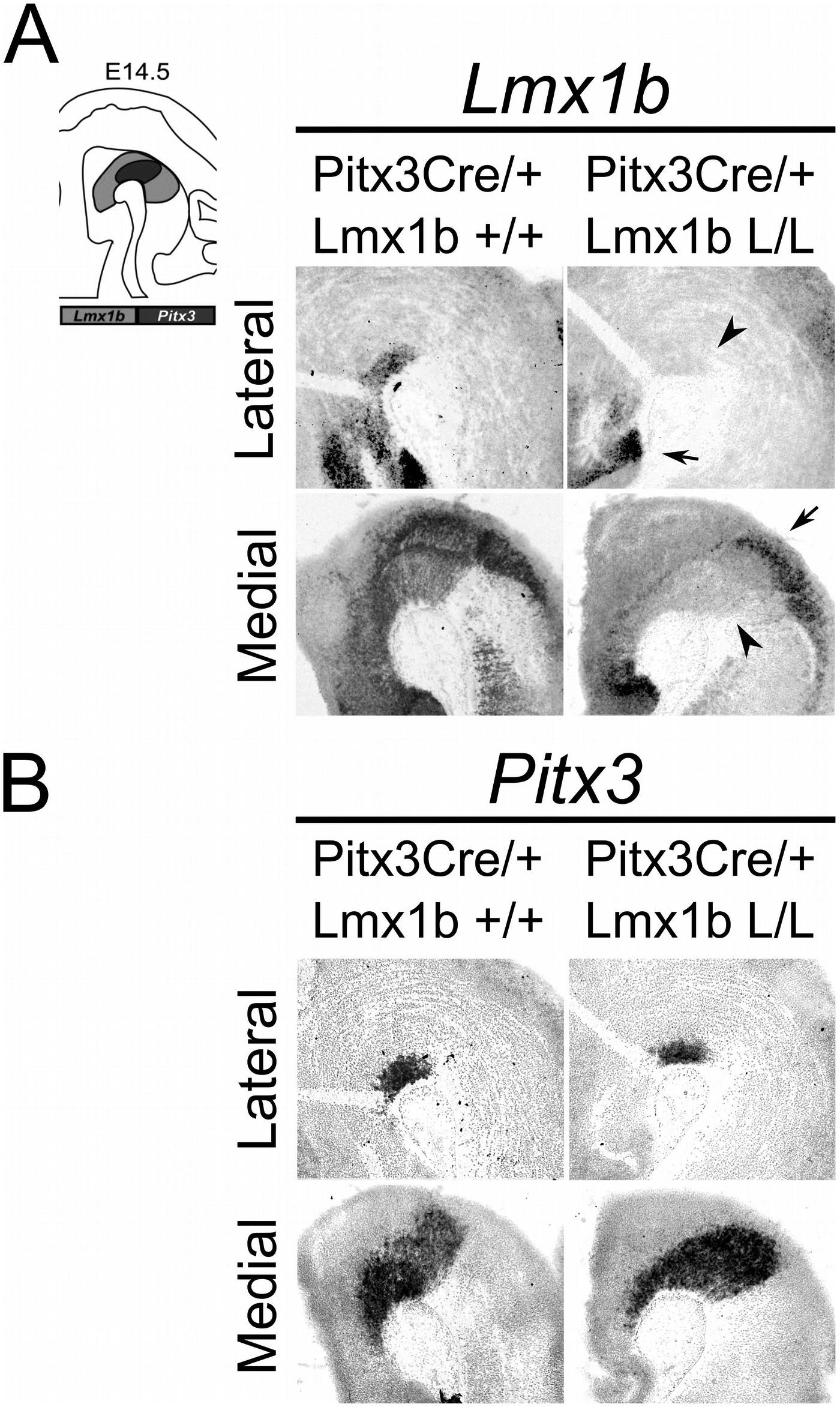
*Lmx1b* expression is specifically lost in the *Pitx3* positive domain of Lmx1b cKO embryos. Analysis of *Lmx1b* and *Pitx3* expression via *in situ* hybridization in E14.5 midbrain sagittal sections. (A) When comparing the *Pitx3Cre*/+; *Lmx1b* L/L to the *Pitx3Cre*/+; *Lmx1b* +/+ a clear loss of *Lmx1b* is observed in the *Pitx3* positive area (arrowheads). However *Lmx1b* expression is maintained in regions that are negative for *Pitx3* (arrows). (B) *Pitx3* expression seems unaffected.

### *Lmx1b* is not essential for early post-mitotic development of mdDA neurons

As described above, the onset of *Pitx3* driven CRE activity is around E13.5 and by E14.5 most genes that define a mature mdDA neuron, like *Th, Vmat2* and *Dat*, are expressed (Arenas et al., 2015; Iversen, 2010). The expression of these genes is dependent on the combined functioning of *Nurr1* and *Pitx3* (Hwang et al., 2003; Jacobs et al., 2009; Munckhof et al., 2003; Nunes et al., 2003; Saucedo-Cardenas et al., 1998; Smidt et al., 2004; Smits et al., 2003). Loss of function studies showed that *Nurr1* mutants fail to induce *Th, Vmat2* and *Dat*, leading to a developmental arrest of mdDA progenitors and cell death (Saucedo-Cardenas et al., 1998; Smits et al., 2003; Zetterström et al., 1997). While *Pitx3* was found to be crucial for the correct specification of mdDA neurons by acting as an activator of the NURR1 transcriptional complex (Jacobs et al., 2009). Since we removed *Lmx1b* in post-mitotic cells, we hypothesized that any effect on the differentiation of mdDA neurons would become apparent in the spatial expression of *Th, Vmat2* and *Dat* at E14.5. In addition to these three factors, we also examined *Aadc*, which is also required for the proper DA neurotransmitter phenotype and is affected in both *Nurr1* and the *Pitx3* mutants, similar to the other marks we examined (Saucedo-Cardenas et al., 1998; Smidt et al., 2004). However this gene is induced in an early state at E10.5 in a *Nurr1*-independent manner (Smits et al., 2003). When analyzing the spatial expression of *Th* via *in situ* hybridization no clear differences could be observed in the lateral or medial sections (Figure 2A). In addition the expression of both *Vmat2* and *Aadc* also seem unaffected by the post-mitotic loss of *Lmx1b* (Figure 2B, C). However when examining the expression of *Dat* (Figure 2D), we observed a different expression pattern between the *Pitx3Cre/+; Lmx1b* +/+ and *Pitx3Cre/+; Lmx1b* L/L embryos. Together these results suggest that post-mitotic deletion of *Lmx1b* does not influence the expression of general DA marks *Th, Vmat2* and *Aadc*, however the expression pattern of *Dat* is altered, suggesting that *Lmx1b* might be involved in defining spatial distribution of *Dat*.

**Figure 2:**
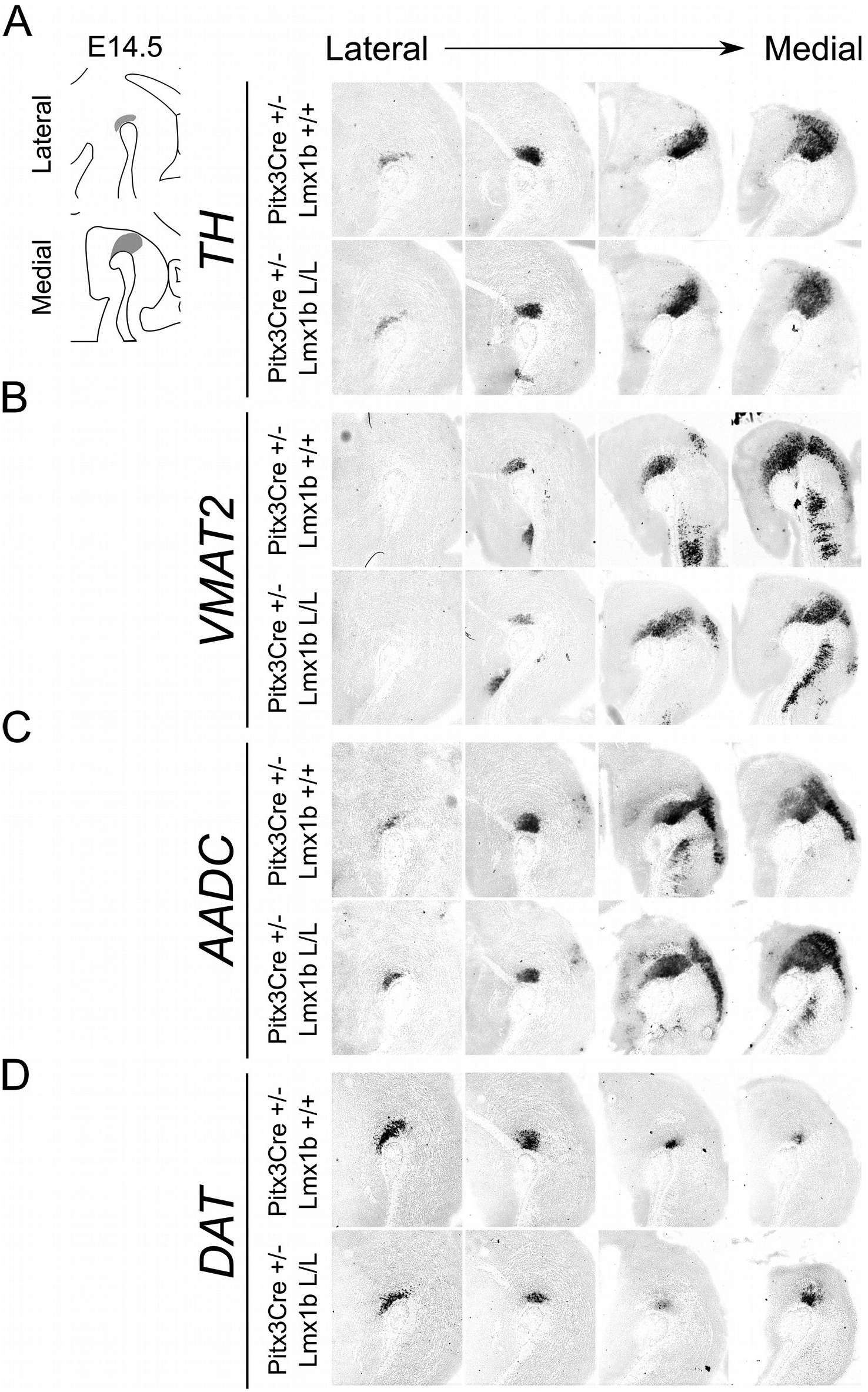
Several genes important for the DA neurotransmitter phenotype do not seem to be affected by the *Pitx3Cre* driven loss of *Lmx1b* at E14.5. (A, B, C, D) *In situ* hybridization of *Th, Vmat2, Aadc* and *Dat* in E14.5 midbrain sagittal sections. (A, B, C) In both lateral and medial sections expression of *Th, Vmat2* and *Aadc* do not seem to be affected in *Pitx3Cre/+; Lmx1b* L/L embryos. (D) However when examining the expression of *Dat* (lowest panel) the pattern seems to be altered in the *Lmx1b* cKO embryos compared to *Pitx3Cre/+; Lmx1b* +/+ embryos.

### Conditional removal of *Lmx1b* results in the loss of TH+ cells in the adult midbrain

Previous studies have already shown that *Lmx1b* expression is continued throughout life (Dai et al., 2008; Laguna et al., 2015; Smidt et al., 2000), suggesting that *Lmx1b* might have a role in neuronal identity, maintenance and survival. A study performed by Doucet-Beaupre *et al.* demonstrated that the combined conditional genetic ablation of *Lmx1a* and *Lmx1b* under the control of the *Dat* promoter causes a degeneration of TH+ cells in 2 months old mice in both the SNc and the VTA (Doucet-Beaupré et al., 2016). To verify whether *Pitx3* driven deletion of *Lmx1b* also influenced neuronal identity and/or maintenance we examined the expression of *Th, Vmat2, Aadc*, and *Dat* in 3 months old midbrains (Figure 3). When comparing *Pitx3Cre*; *Lmx1b* +/+ animals to *Pitx3Cre*; *Lmx1b* L/L animals no obvious alterations in the expression patterns of *Th* and *Vmat2* can be observed (Figure 3A, B), however when analyzing the expression of *Aadc*, alterations can be observed (Figure 3C, arrowsheads) and when comparing *Dat* expression between the wildtype and the *Lmx1b* cKOs an overall reduction can be seen (Figure 3D, arrow heads). To further establish whether changes in expression are caused by the influence of *Lmx1b* on specific gene expression or on neuronal survival in general we aimed to quantify the total amount of TH+ cells in the mdDA population. We performed immunohistochemistry for TH and counted the cells in both the SNc and the VTA (Figure 4A). The total amount of TH+ cells is reduced with ~ 15% (n=3, ** P<0.01, one-tailed) in the cKO compared to wildtype littermates (Figure 4B). The loss of TH+ cells is largest in the VTA (~ 20% loss, ** P<0.01, one-tailed), while ~13% of the cells in the SNc are lost (Figure 4B). Since Laguna *et al.* found a progressive loss of cells between 2 months old cKO mice and 18 months old *DatCre*; *Lmx1a* L/L; *Lmx1b* L/L animals (Laguna et al., 2015), we aimed to investigate whether the reduction in TH+ cells in our model would also further progress. We quantified the amount of TH+ cells in the SNc and the VTA in 12 month old midbrains (Figure5A). A reduction of ~10% of the total TH+ population is observed in *Pitx3Cre*; *Lmx1b* L/L midbrains (n=4, P< 0.05, one-tailed) (Figure 5B). In contrast to 3 months old animals, the loss of TH+ cells is only observed in the SNc, where the amount of TH+ cells is reduced with 15% (n=4, P<0.05, one-tailed) in the cKO midbrains, whereas the amount of TH+ cells in the VTA are not significantly altered (n=4) (Figure 5B). Together these results demonstrate that ablation of *Lmx1b* leads to the loss of TH+ neurons. In the VTA an accelerated loss of cells is observed in the *Lmx1b* cKO, leading to reduced numbers of TH+ cells at 3 months of age. However, over time the number of cells between the wildtype and *Pitx3Cre*/+; *Lmx1b* L/L animals are equalized, suggesting that during aging *Pitx3Cre*/+; *Lmx1b* +/+ animals also lose cells in the VTA. Interestingly, the defect observed in the SNc at 3 months of age, resides with time and is still present at 12 months of age.

**Figure 3:**
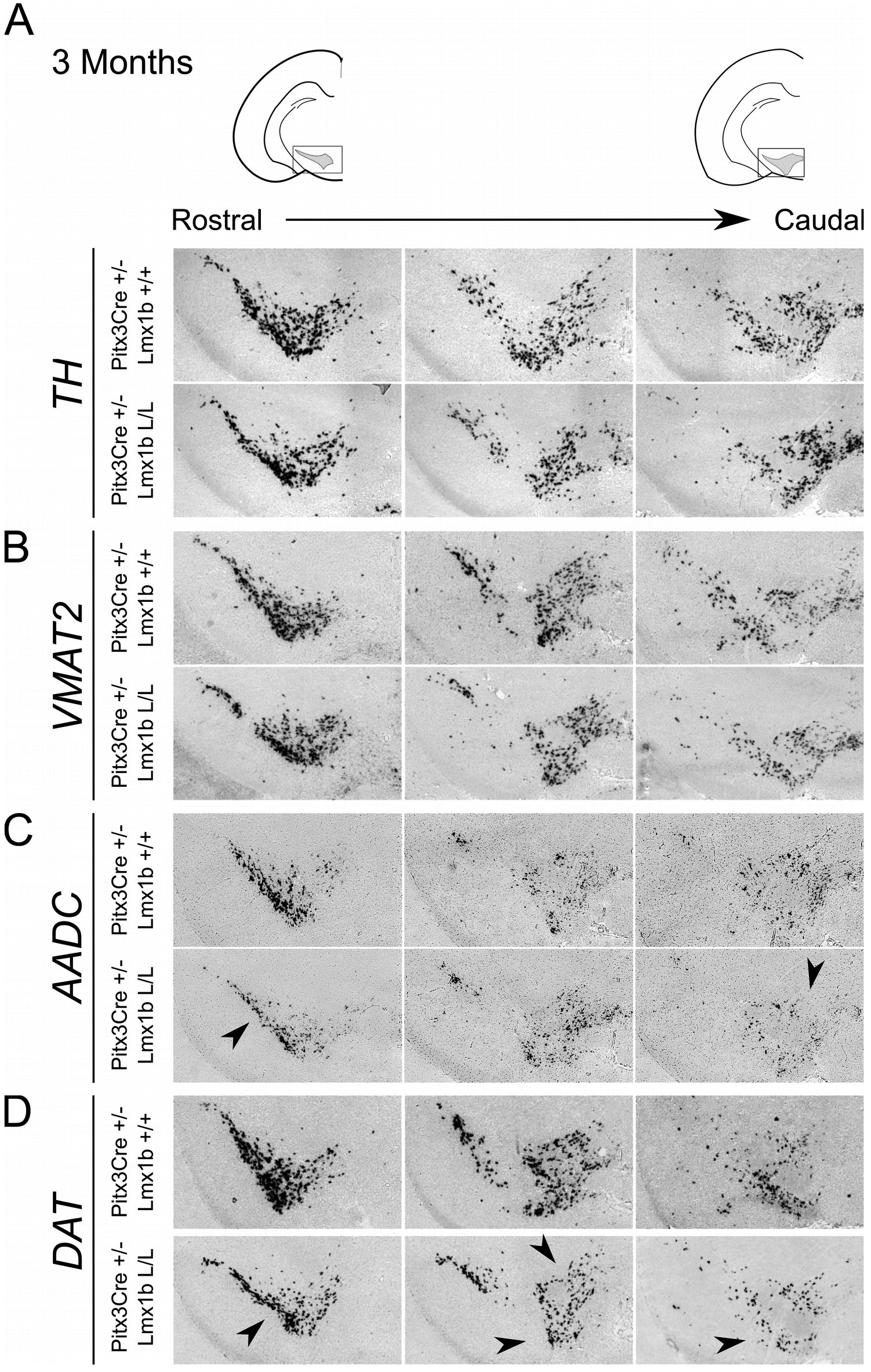
Pitx3 driven deletion of Lmx1b results in a partial loss of *Dat* expression in the adult midbrain. Analysis of *Th, Vmat2, Aadc* and *Dat* in coronal adult section in the *Pitx3Cre*/+; *Lmx1b* L/L mutant via *in situ* hybdrization. (A, B) Expression of *Th* and *Vmat2* seem unaffected in *Pitx3Cre*/+; *Lmx1b* L/L animals, but the *Pitx3* driven deletion of *Lmx1b* results in alteration in the expression of *Aadc* (C, arrowsheads) and an overall reduction in *Dat* expression (D) in both rostral and caudal sections (arrowheads).

**Figure 4:**
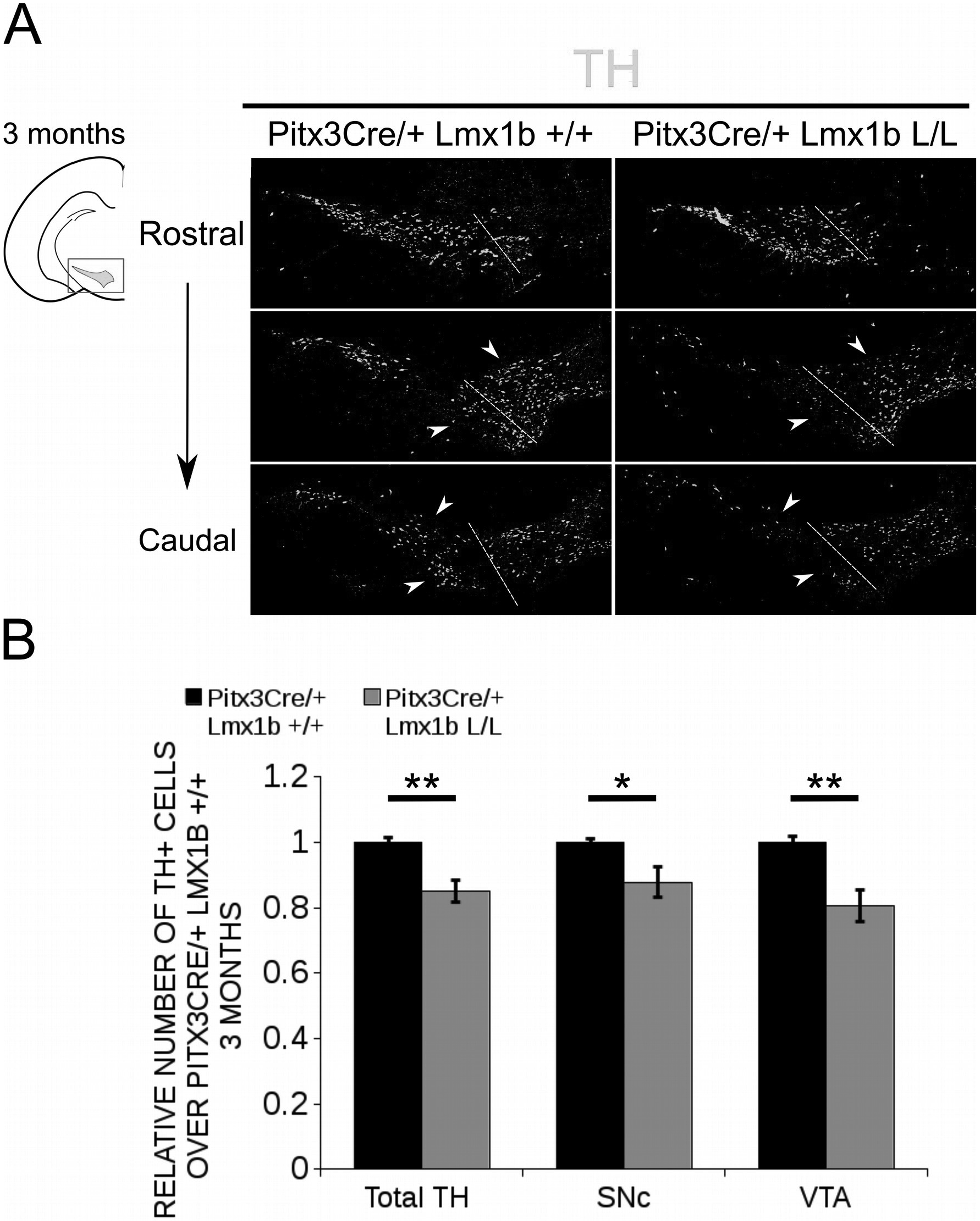
The number of TH+ cells are decreased in Pitx3Cre/+ Lmx1b L/L 3 months old midbrains. (A) Protein expression of TH (green) was evaluated via immunohistochemistry in the adult midbrain of 3 months old *Lmx1b* cKO animals. A loss of TH signal was observed in both the VTA and the SNc (white arrows). The white dotted line represent the border between what is considered SNc and VTA. (b) Quantification of TH+ cells in the adult midbrain of *Pitx3Cre*/+; *Lmx1b* L/L (n=3, grey bars) and *Pitx3Cre*/+; *Lmx1b* +/+ controls (n=3, black bars) shows that the total amount of TH+ neurons is significantly lower (~ 15% loss, ** P< 0.01, one-tailed) and that neurons are lost both in the SNc (~ 13% loss, * P<0.05) and the VTA (~ 20% loss, ** P<0.01, one-tailed). *Pitx3Cre/+ Lmxlb* +/+ animals were set at 1.

**Figure 5:**
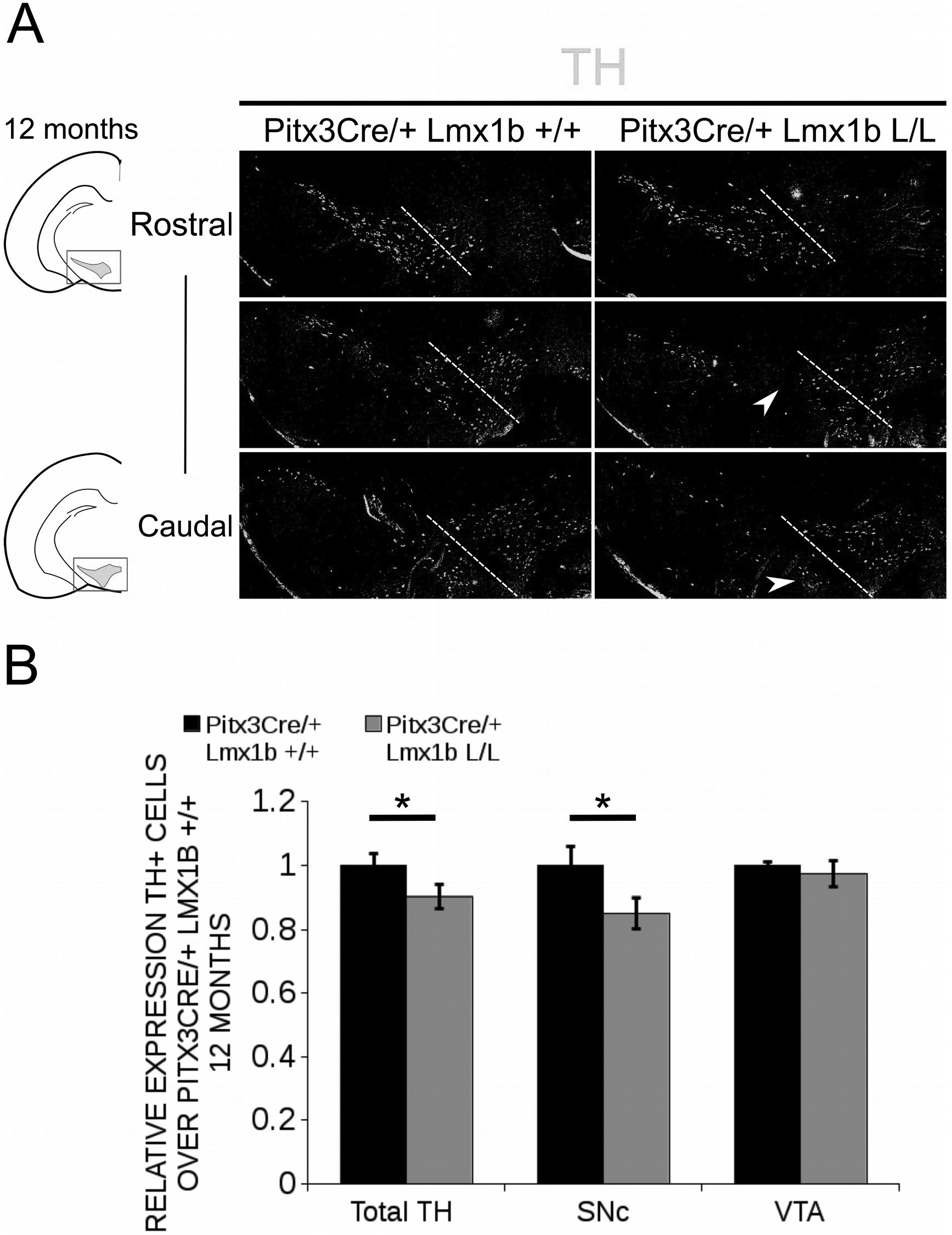
In 12 months old midbrains a reduction in TH+ neurons is observed in Pitx3Cre driven *Lmx1b* cKO animals. (A) Protein expression of TH (green) was evaluated by immunohistochemistry in adult midbrains of 12 months old mutants and wildtypes. A loss of TH signal was observed in the SNc (white arrows). The white dotted line represent the border between what is considered SNc and VTA. (B) Quantification of TH+ cells in the adult midbrains of *Pitx3Cre*/+; *Lmx1b* L/L (n=4, grey bars) and *Pitx3Cre*/+; *Lmx1b* +/+ controls (n=4, black bars) shows that the total amount of TH+ neurons is significantly lower (~ 10% loss, ** P< 0.01, one-tailed) and that neurons are lost in the SNc (~ 15% loss, * P<0.05), but no loss was observed in the VTA (n=4, one-tailed). *Pitx3Cre/+ Lmxlb* +/+ animals were set at 1.

### *Lmx1b* regulates genes involved in neuronal development and acts as a repressor of *Ahd2*

In order to obtain a better insight into which molecular mechanisms are affected during development after the conditional removal of *Lmx1b* we performed next-generation RNA-sequencing on dissected E14.5 midbrains of *Pitx3Cre*/+; *Lmx1b* +/+ and *Pitx3Cre/Lmx1b* L/L embryos (n=3; 2 pooled embryos per biological replicate) (Figure 6A). When using a P-value cut-off of P< 0.05 285 genes were identified, of which 105 were up-regulated and 180 are down-regulated. Among the 20 most regulated genes we found *Aldh1a1 (Ahd2), En1* and *En2* (Figure 6B). *En1* and *En2* have been implicated in mdDA neuronal survival before (Albéri et al., 2004; Simon et al., 2001). It was shown that within animals heterozygous for *En1* lose DA neurons specifically in the SNc (Sonnier et al.,2007). In addition to a role in neuronal survival *En1* has been implicated to play a role in the specification of the mdDA subpopulations. *En1*-deficient embryos demonstrate a down-regulation of *Th, Dat, Vmat2* and *D2R* in rostrolateral sub-population of mdDA neurons, which are destined to become the SNc (Veenvliet et al., 2013). Interestingly, this rostrolateral sub-population is marked by the expression of *Ahd2* (Jacobs et al., 2007; Veenvliet et al., 2013), which shows an upregulation of approximately 2 fold (Figure 6B), suggesting that the loss of *Lmx1b* might specifically affect the rostrolateral subset. To identify other genes associated with mdDA neuronal development and the rostrolateral subset we cross-referenced the list of 285 possible target genes to a list of genes known to be involved in mdDA development (Figure 6C). In addition, we performed an overlay of transcripts of *Lmx1b* target genes with the MAANOVA-FDR analysis of genes regulated by either *En1* (Veenvliet et al., 2013) or *Pitx3* (Jacobs et al., 2011), two genes that have been shown to be essential for the formation of rostrolateral mdDA neurons (Veenvliet et al., 2013). We found 9 genes that were previously associated with mdDA development of which *En1* and *Ahd2* were considered for further study (Figure 6D, red arrows). Furthermore genes regulated by both *Lmx1b* and *En1* were found to be regulated in the same direction (Figure 6E). This in contrast to genes that are regulated by both *Lmx1b* and *Pitx3*, which are mostly regulated reciprocal (Figure 6F), similar to the *Pitx3-En1* interplay (Veenvliet et al., 2013). *En1, En2* and *CD9* were found to be regulated by all three genes, *Lmx1b, En1* and *Pitx3* (Figure 6G), and while they are up-regulated in the *Pitx3* mutant (Figure 6F), they are down-regulated in both *En1* mutants and *Pitx3Cre*/+; *Lmx1b* L/L embryos (Figure 6E). Based on the results from the genome-wide expression analysis and the association of *En1* and Ahd2 with the SNc (Jacobs et al., 2007; Sonnier et al., 2007; Veenvliet et al., 2013), we decided to examine the spatial expression of these two genes. In independent sets, the expression levels were confirmed using qPCR and the spatial expression patterns were studied using *in situ* hybridization. The levels of *En1* were 30% reduced (n=4, *P<0.05, one-tailed) in the *Lmx1b* mutant embryo compared to wildtype (Figure 7A, right panel), validating our RNA-sequencing results. Interestingly, the spatial expression of *En1* is not altered in the *Pitx3Cre*/+; *Lmx1b* L/L embryos compared to *Pitx3Cre*/+; *Lmx1b* +/+ embryos at E14.5 (Figure 7A, left panel), suggesting that *Lmx1b* is important for the expression level of *En1* in all mdDA neurons. When examining the expression pattern of *Ahd2*, an increase in expression can be found in the entire *Ahd2* positive area (Figure 7B) and clear ectopic *Adh2* expression is found in the more medial sections (Figure 7B, left panel, arrowheads). Quantification of the expression levels showed a 2 fold increase in *Adh2* (n=4, ** p<0.01, one-tailed) in the mutant compared to wildtype (Figure 7B, right panel). In summary, we show that *Lmx1b* is important for the regulation of genes associated with mdDA neuronal development, including *En1*, and that *Lmx1b* acts as a repressor of *Ahd2*.

**Figure 6:**
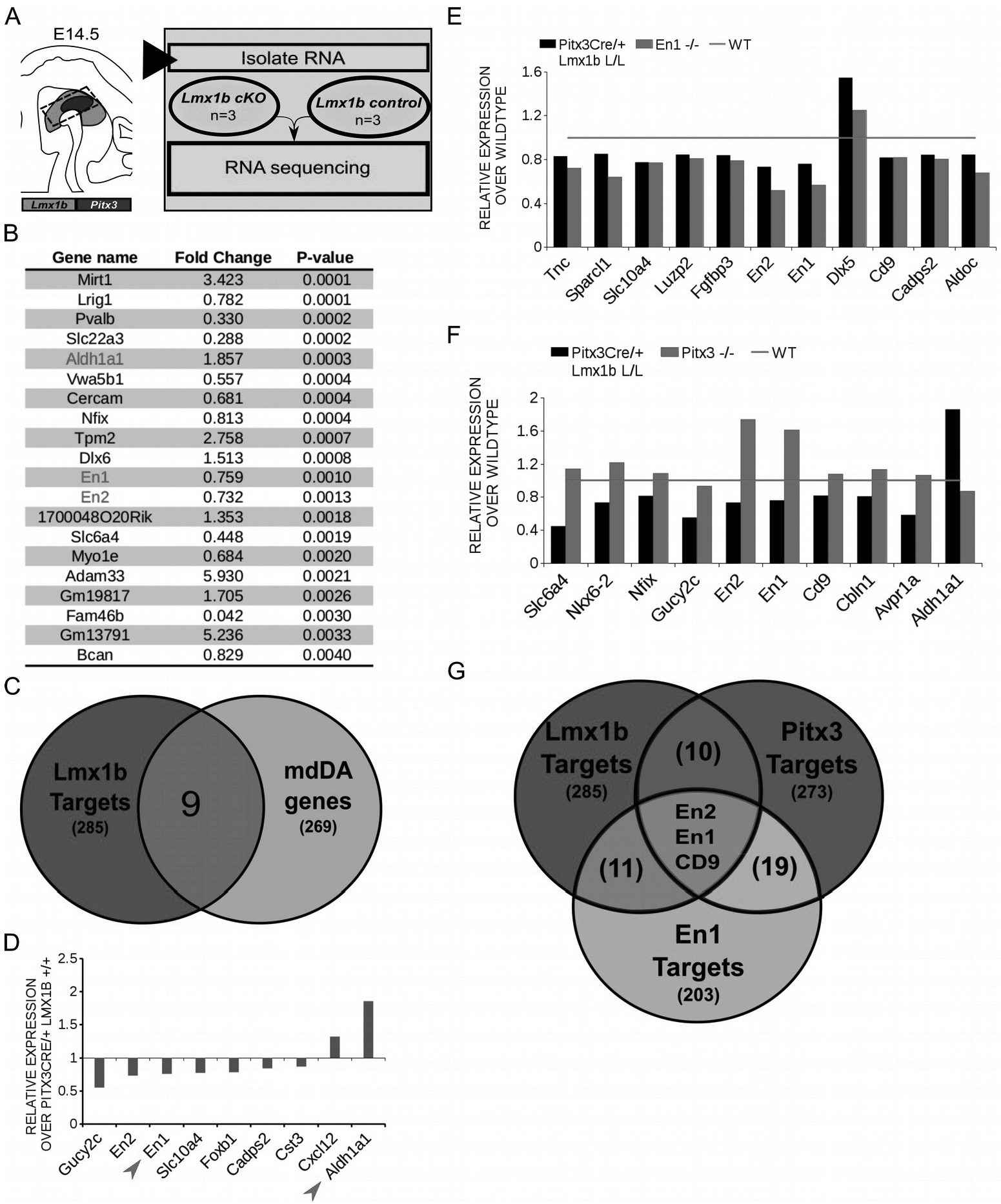
*In vivo* genome-wide expression analysis reveals that *Lmx1b* regulates genes involved in neuronal development, including *En1* and *En2* and works as a repressor of *Ahd2* expression. PANTHER overrepresentation test shows *Lmx1b* is important for nervous system development (red) and for organ development. (B) Among the 20 most regulated genes are *Ahd2* (Aldh1a1, red), which shows an 85% increase, and *En1* (red) and *En2* (red) which are both down-regulated *(En1, ~* 25% loss, *En2*, ~27% lossj (C) Venn diagram illustrating that 9 genes regulated by *Lmx1b* are mdDA enriched. (D) Relative expression levels of the 9 genes that were mdDA enriched. Among the 9 genes *En1* and *Ahd2* were considered most interesting (red arrows). (E) Relative expression of genes regulated by both *En1* and *Lmx1b* demonstrate that genes that are regulated by the loss of *En1* (grey bars) are affected in the same manner in the *Pitx3Cre/+ Lmxlb* L/L E14.5 embryos (black bars). Wildtype was set at 1. (F) When looking at the relative expression of genes regulated by *Pitx3* and *Lmx1b* the opposite effect is observed as with the *En1* mutant. Genes up-regulated in the *Pitx3* mutant are down-regulated in the *Lmx1b* depleted embryos and genes up-regulated in the *Pitx3Cre* driven *Lmx1b* mutant are down- regulated in embryos missing *Pitx3*. (G) Venn diagram showing that *Lmx1b, En1* and *Pitx3* are all involved in regulating *En1, En2* and *Cd9.*

**Figure 7:**
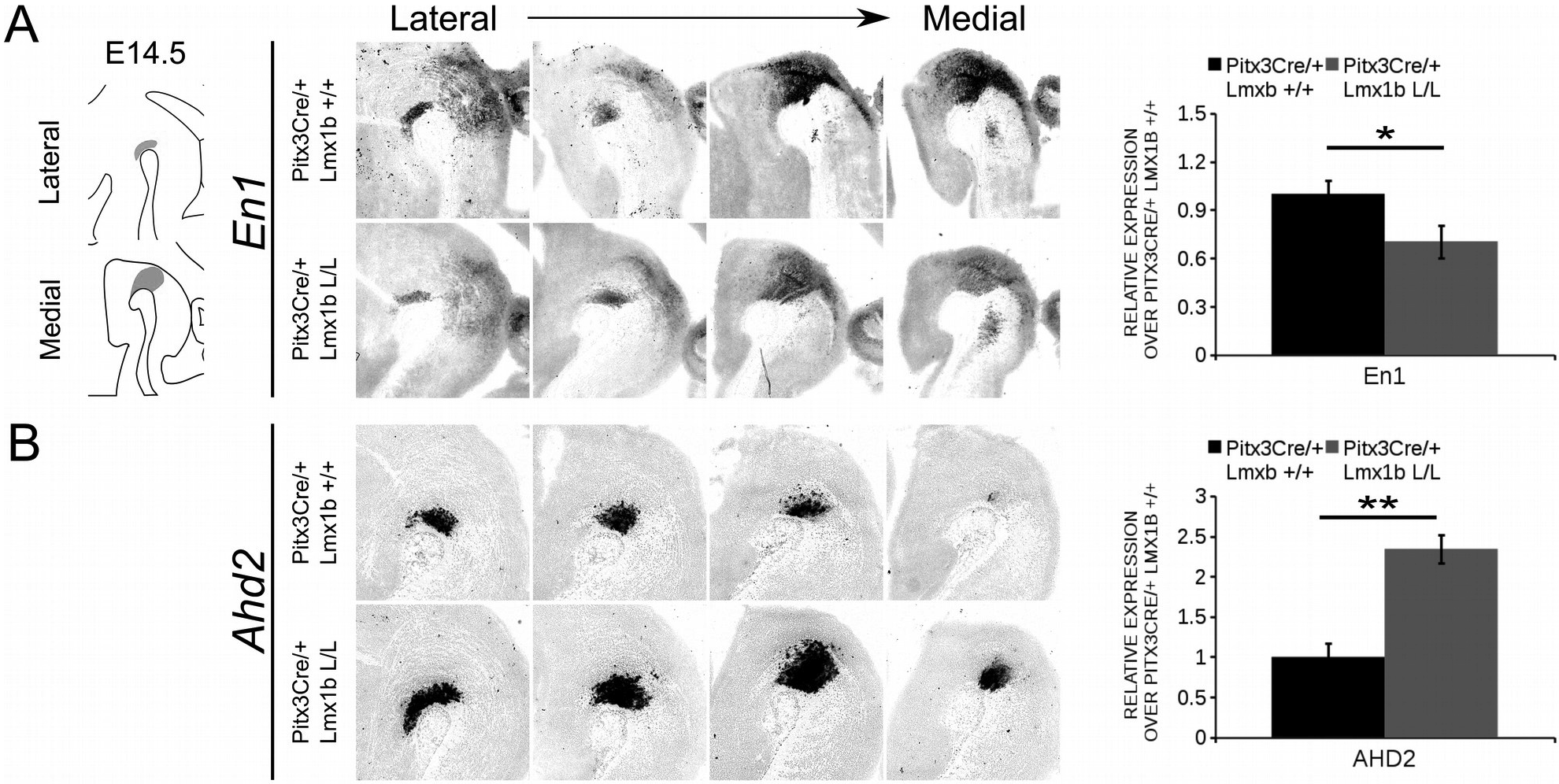
*Ahd2* is up-regulated and ectopically expressed in the *Pitx3Cre* driven *Lmx1b* mutant, together with a generic reduction in *En1* expression in the midbrain of E14.5 mutant embryos. The expression patterns of *En1* and *Ahd2* in E14.5 midbrains of *Lmx1b* cKO and wildtype were analyzed via *in situ* hybridization, followed by qPCR to quantify the mRNA levels. (A) The expression level of *En1* is significantly reduced (~ 30% loss, * p<0.05, one-tailed) in the *Pitx3Cre/+ Lmxlb* L/L E14.5 midbrain (n=4, grey bars) compared to wildtype littermate controls (n=4, black bars). Wildtype levels were set a 1. When visualizing the expression pattern a reduction in *En1* levels can be seen in both lateral and medial sections (green arrows). (B) *Ahd2* expression can be found ectopically in medial section (green arrows) and an up-regulation can be observed in more lateral sections (blue arrows). Quantification of the mRNA levels via qPCR shows an increase of ~ 130% (n=4, ** P<0.01, onetailed) in the mutant (grey bar) compared to the wildtype (black bar). Wildtype was set at 1.

### TH+ cells are lost in the SNc and the VTA of *Pitx3Cre/+ Lmx1b* L/L adult mice

During embryonic development different subsets of mdDA neurons can already be distinguished based on their molecular profile and anatomical position (Smits et al., 2013). As described above, the rostrolateral subset, that will later form most of the SNc, is marked by the expression of *Ahd2* (Jacobs et al., 2011; Veenvliet et al., 2013), while the caudomedial population, destined to become the VTA, expresses *Cck.* (Hökfelt et al., 1980; Veenvliet et al., 2013). It was shown that these subsets are dependent on different transcriptional programs for their proper specification and survival (Jacobs et al., 2011; Kouwenhoven et al., 2017; Panman et al., 2014; Smits et al., 2006; Veenvliet et al., 2013).

The rostrolateral population is dependent on a complex interplay between *Pitx3* and *En1*, in which *En1* is required for the induction of DA phenotype and influences *Pitx3* expression and *Pitx3* is required for antagonizing the caudomedial phenotype by regulating *En1* levels (Veenvliet et al., 2013). When the rostrolateral population is formed, the remaining DA population will obtain a caudomedial phenotype under the influence of *En1* (Bye et al., 2012; Veenvliet et al., 2013). Our model shows lower levels of *En1* and an expansion of the rostrolateral mark *Ahd2* (Figure 7), suggesting that the specification of mdDA subsets is affected. In order to determine whether the loss of *Lmx1b* influences the caudomedial population at E14.5 we examined the expression of *Cck* (Figure 8A). No differences in spatial expression could be observed between the wildtype and *Pitx3Cre*/+; *Lmx1b* L/L embryos when analyzing *Cck* using *in situ* hybridization (Figure 8A, left panel). In addition the levels of *Cck* are not significantly different between the wildtype and the cKO (n=4)(Figure 8A, right panel). Next to analyzing the expression at E14.5, we also decided to examine the spatial expression in 3 month old adult midbrains (Figure 8B). Interestingly, a clear loss of *Cck* expression can be observed in the more lateral positions of the VTA (arrowheads), which seems to match the position where we observed the initial loss in TH+ cells (Figure 4B, arrowheads). To further substantiate the effect of the deletion of *Lmx1b* on the adult mdDA subsets we also examined the expression of *Adh2* in the adult midbrain and quantified the amount of TH+AHD2+ cells in the SNc and the VTA (Figure 9A). The ectopic *Ahd2* expression observed in the embryo could not be detected in the adult stages (Supplemental figure 1) and a ~18% loss of TH+AHD2+ cells (n=3, *P<0.05, one-tailed) is even observed in the SNc (Figure 9B). While the TH+AHD2+ are affected in the SNc, the AHD2 negative population is significantly reduced in the VTA, where ~21% of the TH+AHD2- cells are lost (n=3, **P<0.01, one-tailed) in the mutant compared to *Pitx3Cre/+;Lmx1b* +/+ midbrains (Figure 9C). Together, our data shows that the DA-subsets are differentially affected by the loss of *Lmx1b* in both the embryonic and adult stage.

**Figure 8:**
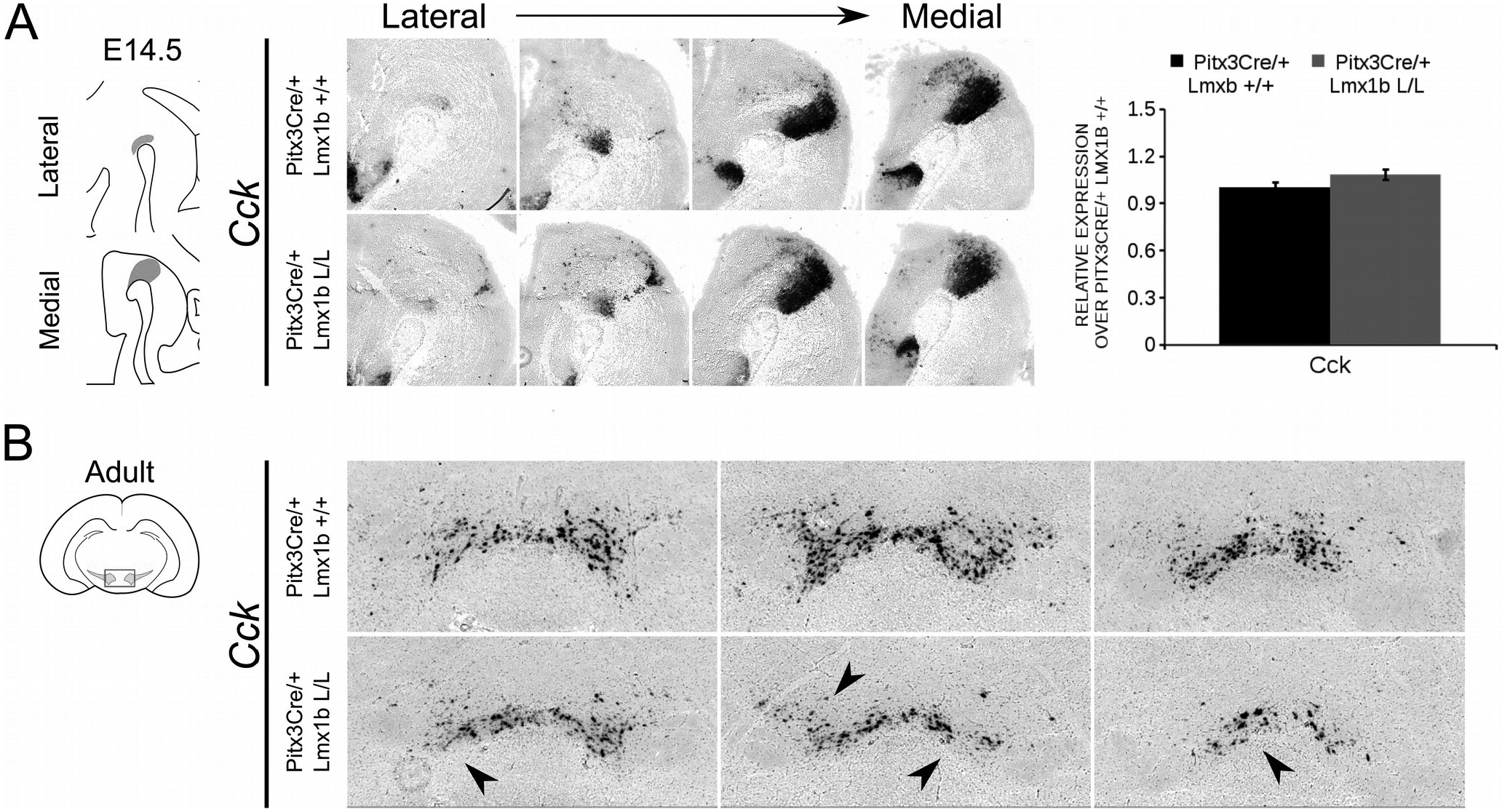
*Cck* expression is not affected in the midbrain of E14.5 *Pitx3cre/+ Lmx1b* L/L embryos, but shows a clear loss in the adult mutant midbrain. (A) Expression of *Cck* was examined using *in situ* hybridization and qPCR. No significant difference in expression level was found between *Lmx1b* conditional mutant E14.5 midbrain (n=4, grey bar) and wildtype littermates midbrains (n=4, black bar). Wildtype levels were set at 1. In addition the expression pattern of *Cck* is comparable between *Pitx3Cre*/+; *Lmx1b* L/L E14.5 midbrain sections and *Pitx3Cre/+ Lmx1b +/+* E14.5 midbrain sections. (B) *In situ* hybridization on the adult midbrain demonstrates a loss of *Cck* expression in the lateral parts of the VTA (arrowheads, n=4 observation).

**Figure 9:**
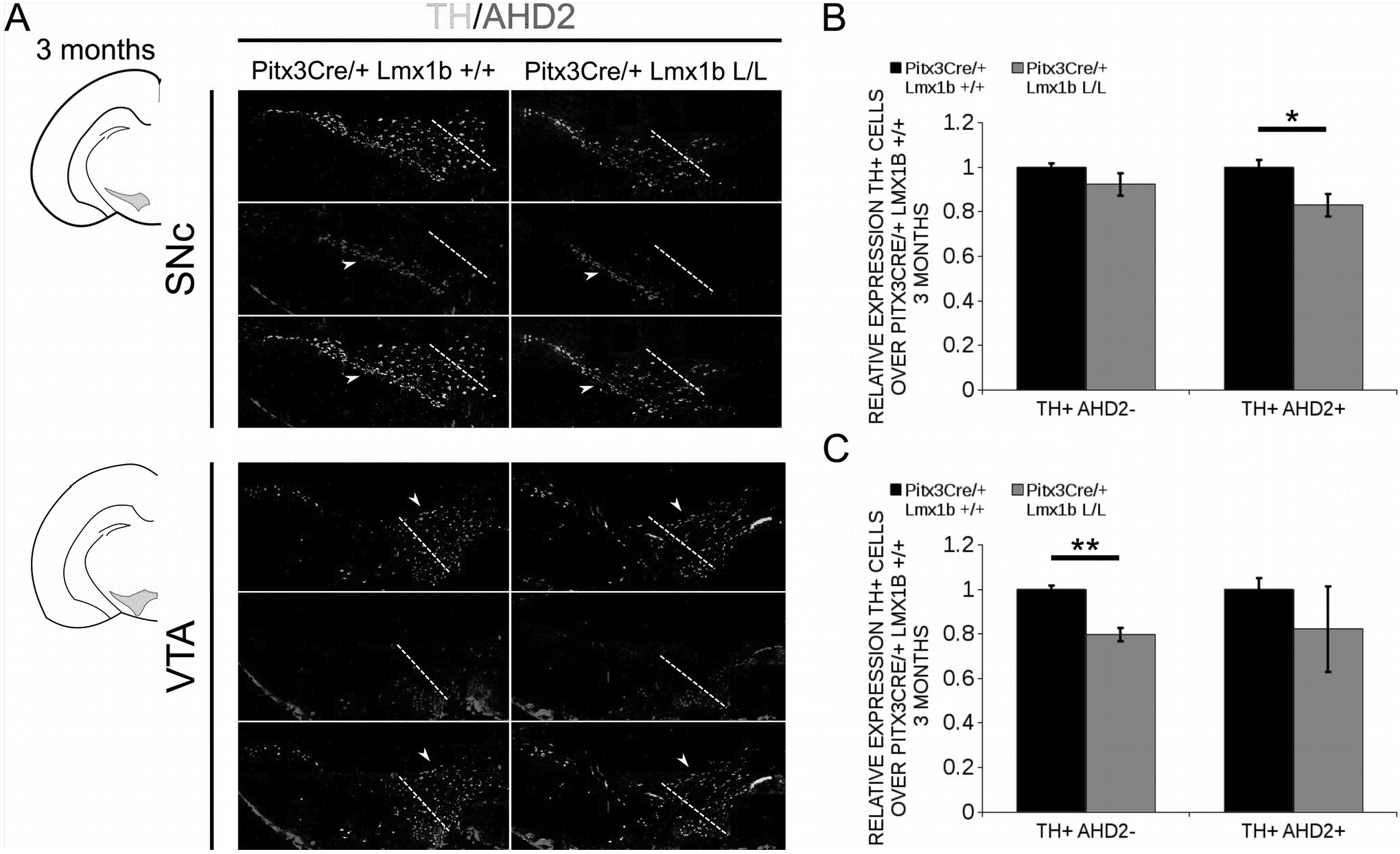
The TH+ cells that are lost in the SNc of 3 months old mice conditionally depleted of *Lmx1b* are also AHD2+, while the TH+AHD2- population is affected in the VTA. Immunohistochemistry of TH (green) and AHD2 (red) in the adult midbrain of 3 months old animals. (A) Assesment of the TH+AHD2+ (yellow) and the TH+AHD2- (green) population in both the SNc and VTA of *Pitx3Cre/+ Lmx1b* L/L animals shows that in the SNc (upper panel) the TH+ AHD2+ neurons are affected, while TH+ neurons affected in the VTA (lower panel) are AHD2- (white arrows). The white dotted line represent the border between what is considered SNc and VTA (B) Quantification of the amount of cells in the SNc shows that the number of TH+AHD2+ neurons are reduced (~18%, n=3, * p<0.05, one-tailed), while the number TH+AHD2- cells are similar to the *Pitx3Cre/+ Lmx1b* +/ + (n=3, black bar). (C) This in contrast to the VTA, where the TH+AHD2+ cells are lost (~ 21% loss, n=3, **P<0.01, one-tailed) in the *Lmx1b* cKO (grey bar) and the TH+AHD2+ cells are not significantly reduced. The number of cells of in the midbrain of *Pitx3Cre*/+; *Lmx1b* +/+ animals were set at 1.

### *Lmx1b* represses *Ahd2* expression independent of *Pitx3*

The transcriptional activation of *Ahd2* in the rostrolateral population has been demonstrated to be dependent on the expression of *Pitx3* (Jacobs et al., 2007). *Pitx3* can interact with a region close to the transcriptional start site of *Ahd2* and loss of expression is observed in *Pitx3* -/- embryos and adult midbrains (Jacobs et al., 2007). As mentioned above, the *Pitx3* driven deletion of *Lmx1b* causes an up-regulation of *Adh2* and to get a better insight into the mechanisms via which *Lmx1b* influences *Ahd2* expression we generated a double mutant for *Pitx3* and *Lmx1b* (*Pitx3CreCre*; *Lmx1b* L/L). We analyzed the expression of *Ahd2, Enl* and *Th* in *Pitx3CreCre*; *Lmx1b* L/L and *Pitx3CreCre*; *Lmx1b* +/+ embryos with *in situ* hybridization (Figure 10A, B, C). When analyzing the expression of *Ahd2* we observed a clear increase in signal in several sections (arrows) in embryos depleted of both *Pitx3* and *Lmx1b* (Figure 10A). In more lateral sections this increase in *Ahd2* was less pronounced (Figure 10A). Importantly, The expression of *En1* and *Th* seem unaffected (Figure 10B, C), suggesting that the ablation of *Lmx1b* only leads to an increase in *Ahd2* expression in a *Pitx3* mutant background. To quantify the mRNA levels we isolated RNA from E14.5 dissected midbrains of *Pitx3CreCre: Lmxlb* L/L and *Pitx3CreCre: Lmx1b* +/+ embryos (Figure 10D). As expected the *Lmx1b* levels were severely reduced as a consequence of the conditional ablation (n=3, *P<0.05, one-tailed). In addition to the loss of *Lmx1b, Ahd2* was up-regulated to ~160% (n=3, *p<0.05, one-tailed), indicating that repression of *Ahd2* by *Lmx1b* does not require *Pitx3*.

**Figure 10:**
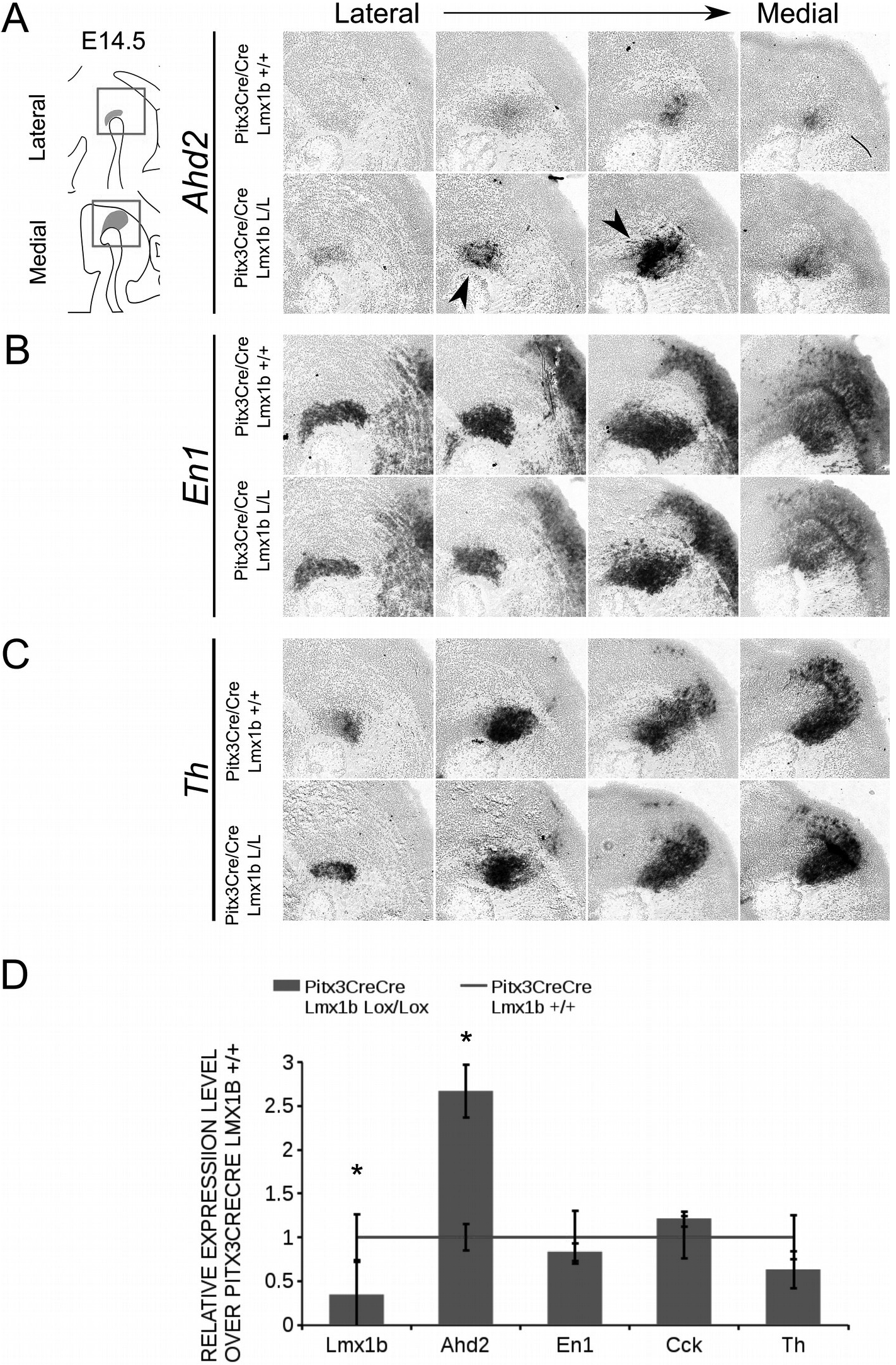
Lmx1b represses Ahd2 independent of Pitx3. The expression patterns of *Adh2, En1* and *Th* were examined using *in situ* hybridization and the expression level *Lmx1b, Ahd2, En1, Cck* and *Th* was verified using qPCR. (A) *Pitx3Cre* driven loss of *Lmx1b* in *Pitx3CreCre* animals caused an increase in *Ahd2* expression in the medial sections of the E14.5 midbrain (green arrows). (B, C) The expression pattern of *En1* did not show any clear differences between *Pitx3CreCre; Lmx1b +/+* and the *Pitx3CreCre Lmx1b* mutant and neither did *Th.* (D) The expression levels of *Lmx1b* were down- regulated with ~76% in conditional mutant (grey bar, n=3, * p<0.05, one-tailed), while *Ahd2* shows an increase of ~160% in the *Pitx3CreCre*; *Lmx1b* L/L (grey bar, n=3, * p~0.05, one-tailed) compared to *Pitx3CreCre*; *Lmx1b* +/+ littermates (red line, n=4). The expression levels of *En1, Cck* and *Th* did not significant changes. Wildtype was set at 1.

AHD2 is an aldehyde dehydrogenase and generates RA and in previous studies it has been shown that a partial recovery of TH+ neurons in the rostrolateral population can be accomplished by maternal supplementation with RA in Pitx3 ablated animals (Jacobs et al., 2007). In addition to the generation of RA, *Ahd2* has also been found to have a protective role for SNc neurons by detoxifying aldehydes (Wey et al., 2012). With partial rescue of *Adh2* expression in the *Pitx3/Lmx1b* double mutant we hypothesized that this might also rescue part of the TH+ cells that are lost in full *Pitx3* mutants in adults. We therefore analyzed the amount of TH+ and TH+AHD2+ cells in the adult midbrain of *Pitx3* mutants and *Pitx3CreCre/Lmx1b* L/L animals by performing immunohistochemistry for TH (green) and AHD2 (red) (Figure 11A). More TH+ cells could be observed in the lateral tier of the SNc (Figure 11A, arrows) and in the VTA (Figure 11A, arrowheads). Quantification showed an ~16% increase in TH+ cells (n=3, ** p< 0.01, one-tailed) and an upward trend for the amount of TH+AHD2+ cells (n=3; p=0.12, one-tailed) in *Pitx3/Lmx1b* double mutant midbrains compared to *Pitx3* mutants (Figure 11B). To summarize our data, by conditionally removing *Lmx1b* in a *Pitx3* mutant background a recovery of *Ahd2* expression is observed at E14.5 and an increase in TH+ cells can be detected in the adult midbrain of *Pitx3CreCre*; *Lmx1b* L/L animals compared to *Pitx3* mutants alone.

**Figure 11:**
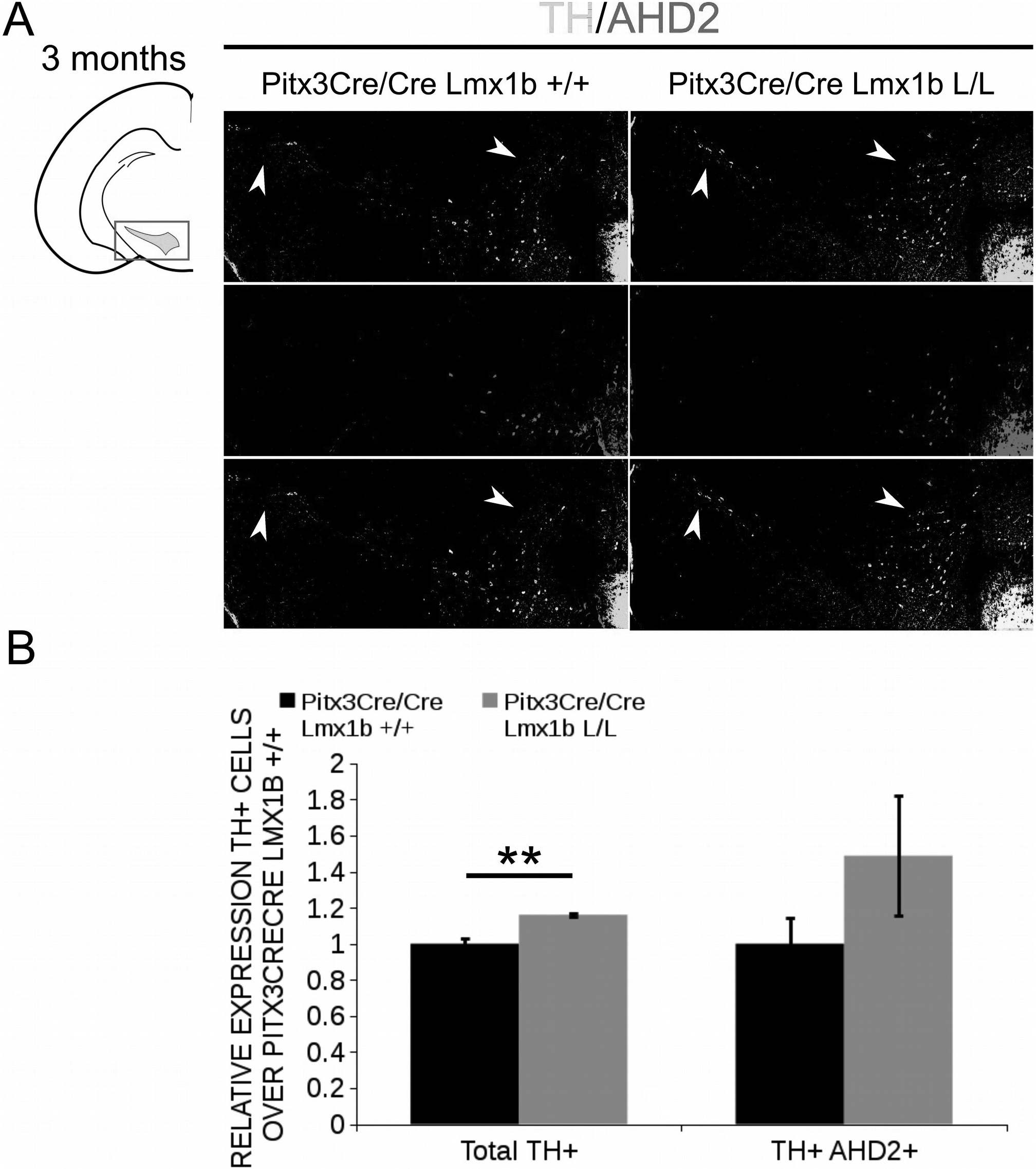
Additional Lmx1b deletion partially rescues the cell loss observed in Pitx3 mutants alone. TH (green) and AHD2 (red) expression was assessed using immunohistochemistry in coronal midbrain sections of 3 months old animals. (A) More TH+ cells were observed in lateral tier of the SNc (white arrows) and in the VTA (white arrowhead). The white dotted line represent the border between what is considered SNc and VTA. (B) Quantification of the total amount of TH+ cells shows an increase in TH+ cells of ~16% (n=3, ** P< 0.01, one-tailed). The number of TH+AHD2+ cells are not significant increased, but shows an upward trend (n=3, P= 0.12, one-tailed). *Pitx3CreCre*; *Lmx1b* +/+ was set at 1.

## Discussion

*Lmx1b* has been associated with several developmental processes (Chen et al., 1998; Adams et al.,2000; Smidt et al., 2000; Guo et al., 2007; Deng et al., 2011; Yan et al., 2011). During early development *Lmx1b* is a critical component of the positive feedback loop that maintains genes associated with IsO functioning. When studying *Lmx1b* null mutants it was demonstrated that *Wnt1, Fgf8, Enl* and *En2* all require *Lmx1b* to maintain their expression at the IsO and in mdDA progenitors (Adams et al., 2000; Guo et al., 2007; Anderegg et al., 2013; Sherf et al., 2015). The loss of *Lmx1b* initially seems to predominantly affect the lateral positioned mdDA progenitor pool, however the medial located progenitors are arrested in their development as they fail to co-express *Th* and *Pitx3* and are lost during later stages (Deng et al., 2011; Smidt et al., 2000). Next to a developmental role a recent study also linked *Lmx1b* to cellular homeostasis of mdDA neurons in adult mice (Laguna et al., 2015). By using a *Dat* driven Cre, *Lmx1b* was removed in postmitotic mdDA neurons, which led to dysregulation of the autophagic-lysosal pathway, abnormal synaptic responses and reduced levels of striatal TH and DAT expression, all processes that contribute to a Parkinson’s disease pathology (Laguna et al., 2015). Here we showed that *Lmx1b* has a variety of functions during postmitotic development, including subset specification and neuronal survival. In line with its function in early development, we found that a loss of *Lmx1b* in postmitotic neurons leads to reduced levels of *En1* and *En2* at E14.5. Both of these factors have be associated with the survival of mdDA neurons (Albéri et al., 2004; Simon et al., 2001) and *En1* +/- animals show a progressive loss of TH+ cells in the ventral midbrain from 3 weeks of age onward (Sonnier et al., 2007). The lower levels of *En1* might contribute to the observed reduction in the amount of TH+ cells in 3 month old *Pitx3Cre*/+; *Lmx1b* L/L animals, however in heterozygous *En1* mice the SNc is mostly affected, while in our model initially the VTA demonstrates the biggest loss of TH+ cells. Notably, after one year the defect resides in the SNc suggesting a change in timing of neuronal survival which culminates in loss of SNc neurons after 12 months in line with En1 mutant animals. In addition to a role in neuronal survival we also found a role for *Lmx1b* in subset specification. *Pitx3* driven deletion of *Lmx1b* led to a significant increase in the rostrolateral mark *Ahd2* in E14.5 embryonic midbrains. In addition, *Ahd2* expression was found ectopically in medial sections of the ventral mesencephalon, overlapping with the *Cck* positive domain. A corresponding shift in the distribution of *Dat* expression was observed, favoring the VTA over the SNc. However, the shift in rostrolateral marks towards the more medial DA domain did not affect the expression of *Cck* during development. The transcriptional activation of *Ahd2* has been demonstrated to be dependent on the expression of *Pitx3* and *En1* (Jacobs et al., 2007; Veenvliet et al., 2013). *Pitx3* can interact with a region close to the transcriptional start site of *Ahd2* and loss of expression is observed in *Pitx3* -/- embryos and adult midbrains (Jacobs et al., 2007). The ablation of *Lmx1b* did not influence the expression levels of *Pitx3* itself (not shown) and also the spatial expression of *Pitx3* was not affected.

In addition, spatial expression of *Ahd2* was partially recovered in *Pitx3/Lmx1b* double mutants compared to *Pitx3* mutants, suggesting that the regulation of *Ahd2* by *Lmx1b* is a PITX3-independent process. Interestingly, the *Ahd2* promoter contains several conserved FLAT-elements, to which LMX1B can bind directly (Rascle et al., 2009), suggesting that LMX1B might directly influence *Ahd2* expression. AHD2 is involved in the generation of retinoic acid (RA) from retinol and Jacobs *et al.* demonstrated that administration of RA to the maternal diet of *Pitx3* -/- animals led to an increase in TH+ cells during development and an increased innervation of the dorsal Striatum (Jacobs et al., 2007). Although we only observed a partial rescue of *Ahd2* expression, we did observe an increase in TH+ cells in the midbrain of adult *Pitx3CreCre*; *Lmx1b* L/L animals.

Overall, our data shows that *Lmx1b* is important for the expression level of survival factors *En1* and *En2* during development and for the survival of a specific group of mdDA neurons postnatally. Next to a role in survival we found that *Lmx1b* functions as a transcriptional repressor of *Ahd2* independent of Pitx3. This underscores the presence of developmental programs that lead to mdDA subsets and that *Lmx1b* is part of the complex network of interacting transcription factors that specify these subsets.

## Acknowledgements

We would like to thank Dr. Ralph Witzgall of the university of Regensburg (Germany) for the kind gift of the Lmxlb-floxed mouse

---

**Supplemental Figure 1:**
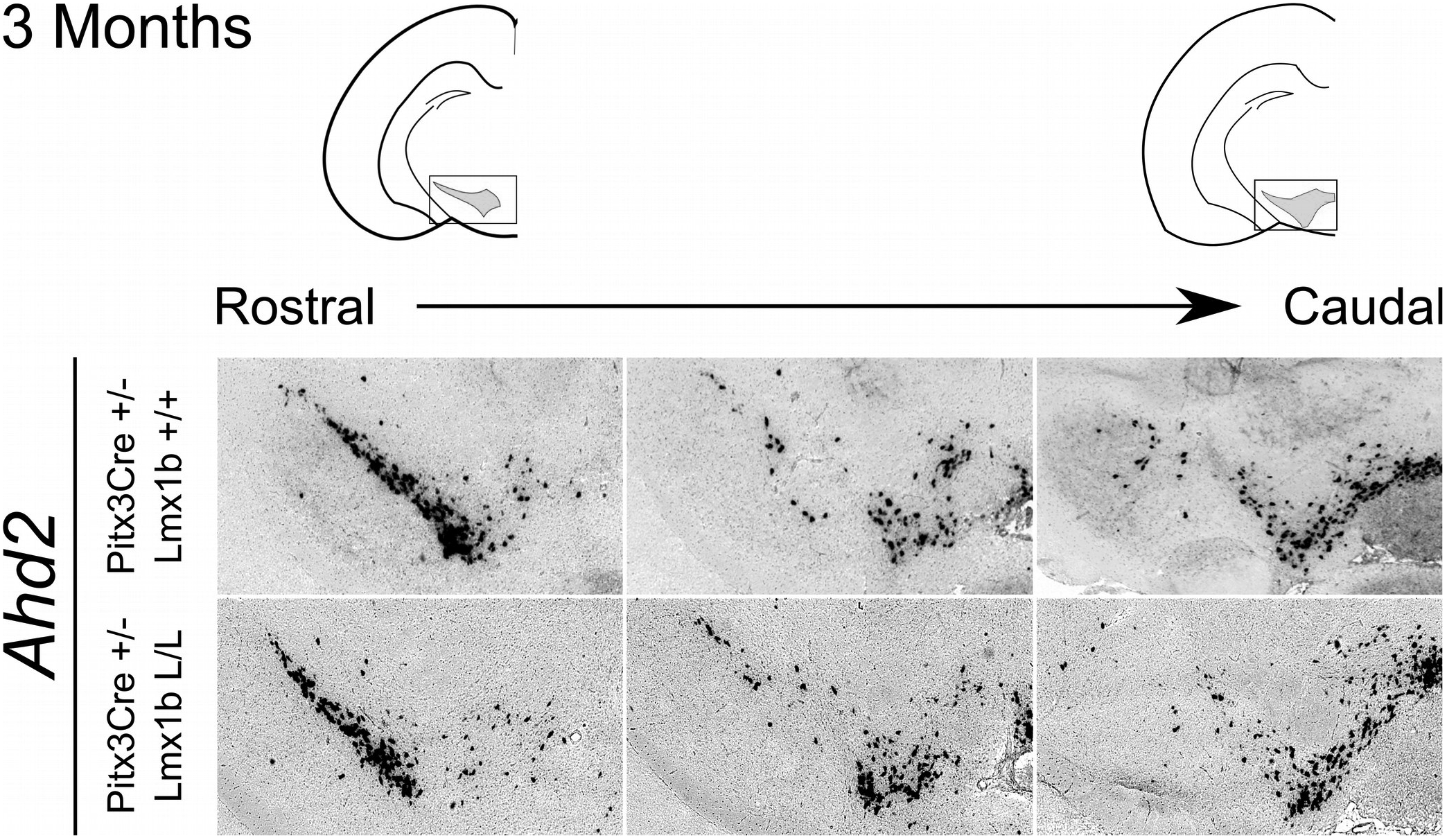
No differences could be detected in the expression pattern of *Ahd2* between *Pitx3Cre*/+; *Lmx1b* +/+ 3 months old animals and *Pitx3Cre*/+; *Lmx1b* L/L animals. Analysis of *Ahd2* in coronal adult section in the *Pitx3Cre*/+; *Lmx1b* L/L mutant via *in situ* hybridization. Expression of *Ahd2* seems similar in *Pitx3Cre*/+; *Lmx1b* L/L animals compared to wildtype.

